# Unveiling the enigma of Brain-resident immune cells

**DOI:** 10.1101/2023.09.26.559602

**Authors:** Sunsook Hwang, Juneil Jang, Kyungsoo Park, Yeong Shin Yim

## Abstract

The immune system has been extensively studied in traditional immune hubs like the spleen and lymph nodes. However, recent advances in immunology highlight unique immune cell characteristics across anatomical compartments. In this study, we challenged conventional thinking by uncovering distinct immune cell populations within the brain parenchyma, separate from those in the blood, meninges, and choroid plexus, with unique transcriptional profiles. Brain-resident immune cells are not derived from maternal immune cells, and age-related changes, with an increase in CD8^+^ T cells in aged mice, are noted. Alzheimer’s disease (AD) alters microglia’s interaction with brain-resident immune cells, emphasizing immune-brain dynamics. Furthermore, we reveal dynamic immune cell interactions and essential cytokine roles in brain homeostasis, with stable cytokine expression but emerging signaling pathways in AD. In summary, this study advances our understanding of brain-resident immune cells in both normal and pathological conditions.

## Introduction

The immune system serves as a sentinel for detecting pathogen invasions, tissue damage, malignancies, wounded healing, and tissue remodeling, conducting an indispensable role in maintaining overall health. To effectively fulfill its multifaceted functions, the immune system is strategically distributed throughout the body, with a functional presence in every tissue and organ. Historically, research on the immune system has primarily focused on cells derived from accessible immune hubs such as the spleen and lymph nodes, due to the logistical challenges associated with obtaining sufficient cell numbers from other anatomical sites. However, recent years have witnessed a paradigm shift in our understanding of immune cell biology, as we now recognize that immune cells exhibit distinct phenotypes, differentiation states, longevity, turnover rates, functions, and regulatory mechanisms based on their anatomical compartment of residence^1,2^.

Among the various tissues in the body, the brain has long remained an underexplored organ in terms of immune interactions, particularly under steady-state conditions. The prevailing concept of brain immune privilege posited that the brain, protected by the blood-brain barrier, was largely insulated from immune influence, with microglia being the sole recognized immune cells within this intricately organized organ^3–5^. However, over the past two decades, groundbreaking research has illuminated the indispensable role of neuroimmune communication in sustaining brain homeostasis and function, extending well beyond pathological scenarios^6^. While in a homeostatic state, most immune cells are primarily localized within the meninges and choroid plexus, with only a sparse population residing within the brain parenchyma, their significance in this context has often been overlooked^7^. Nonetheless, emerging studies have begun to shed light on the substantial impact of immune cells in the brain parenchyma on brain functions such as social behavior, learning, and memory^8,9^.

This paradigm shift necessitates a comprehensive investigation into the tissue-resident immune cells within the brain, their unique characteristics, functions, and contributions to neuronal functions. In this study, we reveal the presence of unique immune cell populations in the brain parenchyma, distinct from those in the blood, meninges, and choroid plexus. We observed age-related shifts in immune composition, with CD8^+^ T cells prevailing in aged mice. Moreover, these immune cells are evenly distributed throughout the brain and originate from offspring rather than maternal sources. Our findings also emphasize the intricate interplay between microglia and brain-resident immune cells, altered in a pathological condition, Alzheimer’s disease, underscoring the pivotal role of immune-brain interactions in maintaining brain homeostasis.

## Results

### The brain has immune cells like other organs

Recent studies have indicated the presence of immune cells in the brain, while it is still controversial^8–12^. The main concerns about this new finding are mainly from 2 parts. 1) Are these immune cells purely from brain parenchyma? Not from the blood contamination? 2) Is the mouse in a non-pathological condition? To eliminate the first concern, we established a protocol to isolate brain-specific immune cells. Due to the abundant distribution of small blood vessels in the brain, we employed an experimental technique to separate immune cells present in brain tissue from blood-derived immune cells^13^. We stained circulating immune cells with anti-CD45 antibody before obtaining brain tissue from mice, allowing us to discriminate immune cells present in brain tissue from those in the blood. After three minutes, mice were perfused with cold PBS, and the brain tissues were harvested. Meninges, choroid plexus, and cerebrospinal fluid (CSF) were further separated. Subsequently, we enriched immune cells with Percoll gradient centrifuge after tissue digestion with Collagenase IV and DNaseI. Samples were subjected to Percoll gradient centrifugation to remove debris and myelin. Enriched immune cells were then FACS analyzed using a set of immune cell markers (Table 1, Fig 1a, Extended Data Fig 1). A variety of immune cells that clearly distinguished from blood-derived immune cells were observed. Brain tissue-specific immune cells are significantly different in the composition of circulating blood-derived immune cells (Fig 1b and c). Surprisingly, Brain parenchyma exhibits different constituent components compared to the meninges and the choroid plexus. These data showed that our method effectively isolates immune cells in the brain, revealing a distinct composition compared to blood-derived immune cells. Furthermore, no indications of previous inflammation were observed when examining the proportion of neutrophils and naive CD4+ and CD8+ T cells (Extended Data Fig 2)^14^.

**Table 1.**
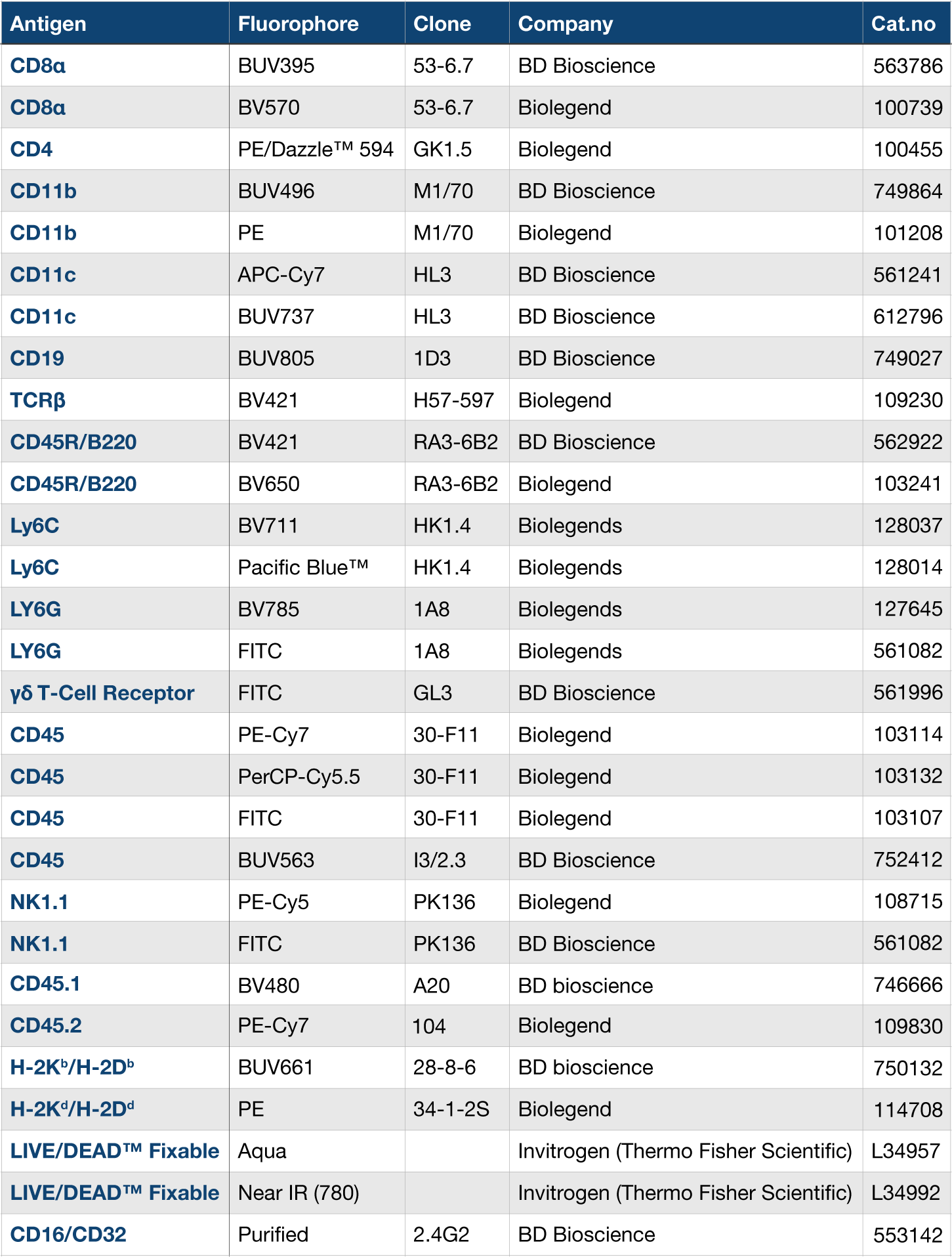
The antibody list for immune cell profiling in mouse brain.

| Antigen | Fluorophore | Clone | Company | Cat.no |
| --- | --- | --- | --- | --- |
| CD8α | BUV395 | 53-6.7 | BD Bioscience | 563786 |
| CD8α | BV570 | 53-6.7 | Biolegend | 100739 |
| CD4 | PE/Dazzle™ 594 | GK1.5 | Biolegend | 100455 |
| CD11b | BUV496 | M1/70 | BD Bioscience | 749864 |
| CD11b | PE | M1/70 | Biolegend | 101208 |
| CD11c | APC-Cy7 | HL3 | BD Bioscience | 561241 |
| CD11c | BUV737 | HL3 | BD Bioscience | 612796 |
| CD19 | BUV805 | 1D3 | BD Bioscience | 749027 |
| TCRβ | BV421 | H57-597 | Biolegend | 109230 |
| CD45R/B220 | BV421 | RA3-6B2 | BD Bioscience | 562922 |
| CD45R/B220 | BV650 | RA3-6B2 | Biolegend | 103241 |
| Ly6C | BV711 | HK1.4 | Biolegends | 128037 |
| Ly6C | Pacific Blue™ | HK1.4 | Biolegends | 128014 |
| LY6G | BV785 | 1A8 | Biolegends | 127645 |
| LY6G | FITC | 1A8 | Biolegends | 561082 |
| γδ T-Cell Receptor | FITC | GL3 | BD Bioscience | 561996 |
| CD45 | PE-Cy7 | 30-F11 | Biolegend | 103114 |
| CD45 | PerCP-Cy5.5 | 30-F11 | Biolegend | 103132 |
| CD45 | FITC | 30-F11 | Biolegend | 103107 |
| CD45 | BUV563 | I3/2.3 | BD Bioscience | 752412 |
| NK1.1 | PE-Cy5 | PK136 | Biolegend | 108715 |
| NK1.1 | FITC | PK136 | BD Bioscience | 561082 |
| CD45.1 | BV480 | A20 | BD bioscience | 746666 |
| CD45.2 | PE-Cy7 | 104 | Biolegend | 109830 |
| H-2K<sup>b</sup>/H-2D<sup>b</sup> | BUV661 | 28-8-6 | BD bioscience | 750132 |
| H-2K<sup>d</sup>/H-2D<sup>d</sup> | PE | 34-1-2S | Biolegend | 114708 |
| LIVE/DEAD™ Fixable | Aqua |  | Invitrogen (Thermo Fisher Scientific) | L34957 |
| LIVE/DEAD™ Fixable | Near IR (780) |  | Invitrogen (Thermo Fisher Scientific) | L34992 |
| CD16/CD32 | Purified | 2.4G2 | BD Bioscience | 553142 |

**Figure 1.**
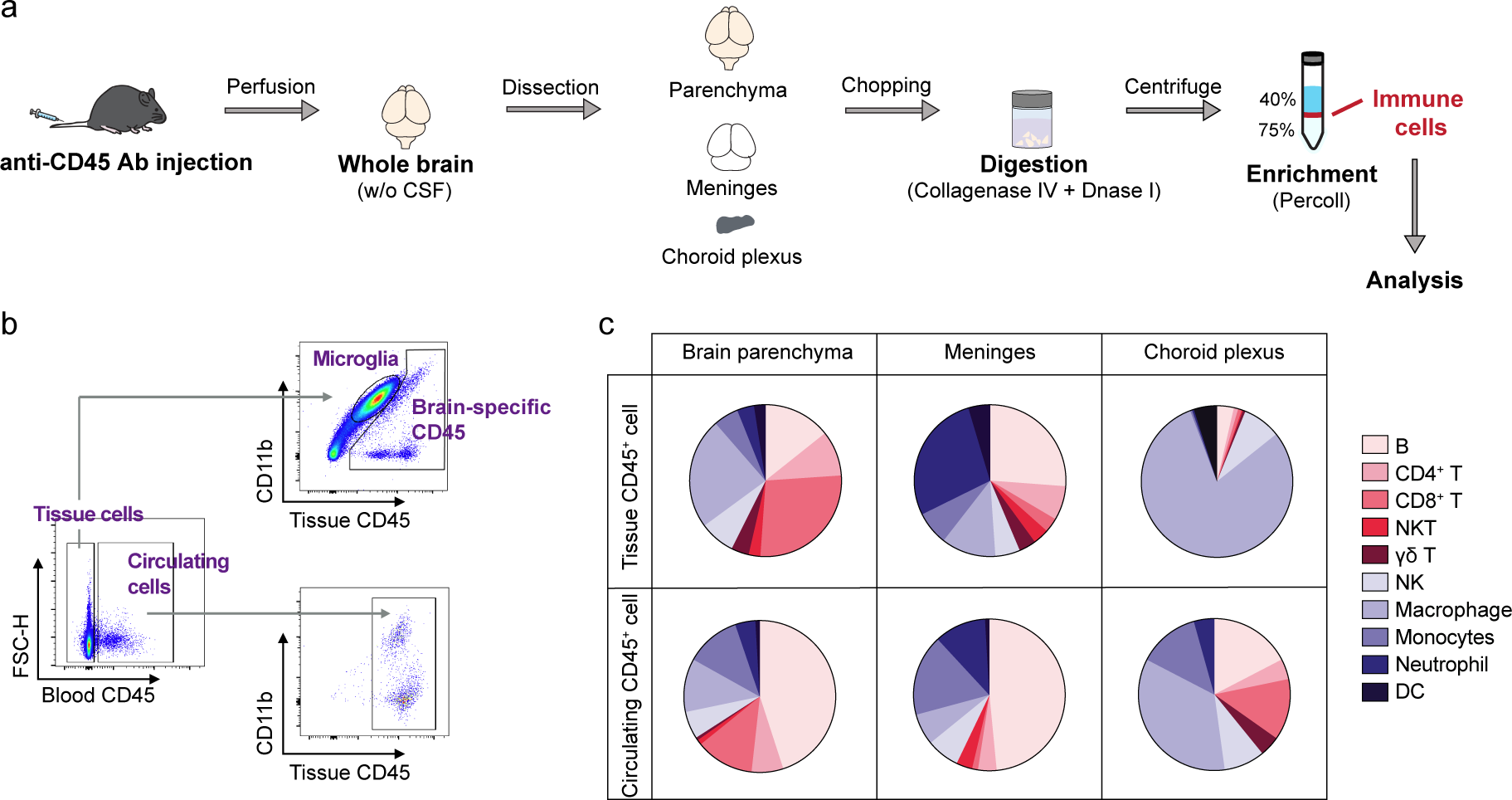
The immune cell composition of the brain parenchyma differs from that of the meninges and choroid plexus. (**a**) Schematic diagram of the experimental method for isolation of brain resident immune cells. Mice received an anti-CD45 antibody injection prior to perfusion. The whole brain was then divided into parenchyma, meninges, and choroid plexus, and the enriched immune cells underwent subsequent analysis. (**b**) The representative FACS plots illustrate the strategy for distinguishing tissue-resident immune cells from circulating blood immune cells using anti-CD45. (**c**) Pie plot indicating the distribution of tissue-resident or circulating blood immune cell profile in the brain parenchyma, meninges, and choroid plexus (n = 4 [brain parenchyma]; n = 6 [meninges]; n = 4 [choroid plexus] from 4 independent experiments).

Next, we analyzed brain parenchymal immune cells in depth to see the age and sex effects on the immune cell composition. When examining total CD45^+^ immune cells in both male and female mice, we observed a significant increase in the number of immune cells in brain parenchyma for mice aged 24 months or older (Fig 2a and b). We also examined the distribution of specific immune cell types based on sex and age. In 4-week-old males, B cells and macrophages were the most abundant, while in females, macrophages predominated over other cell types (Fig 2c and d). Adult mice showed an increased proportion of CD8^+^ T cells in both males and females (Fig 2e and f), with CD8^+^ T cells being the most prevalent in aged mice (Fig 2g-i). Moreover, we conducted an examination to determine if immune cells exhibited distinct localization within specific brain regions. To achieve this, we divided the brain into eight different subregions based on their anatomical structures, encompassing the olfactory bulb, cerebral cortex, hippocampus, hypothalamus, cerebellum, midbrain, hindbrain, and striatum+thalamus. We also included meningeal immune cells as a control. Unexpectedly, our analysis revealed that in both 4-week-old and adult mice, none of the ten immune cell types exhibited regional specificity within the brain (Fig 3). Together, the findings indicate that the brain parenchyma has its own immune system like other non-lymphoid organs and it can be separated from meninges, choroid plexus, and CSF.

**Figure 2.**
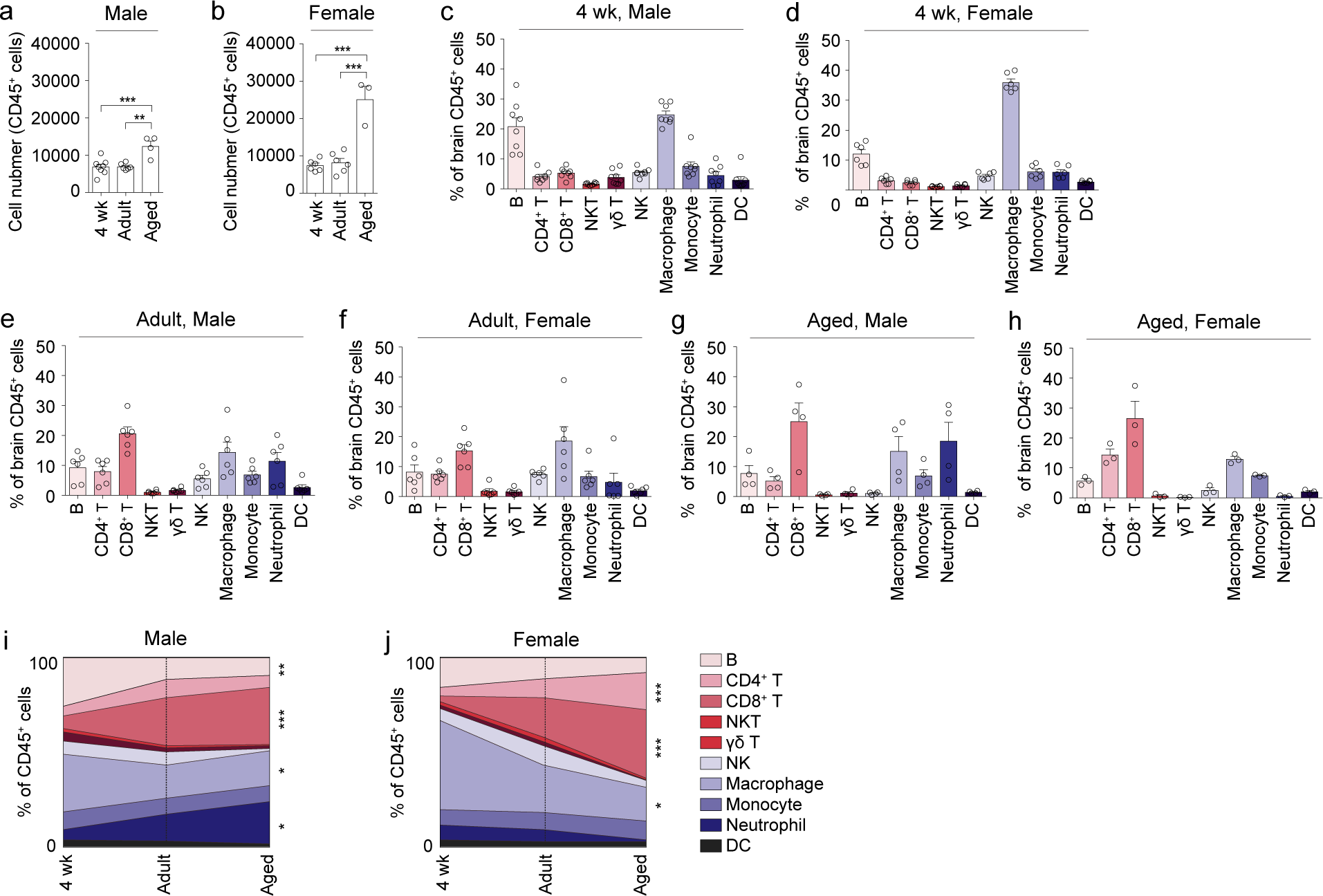
Immune cell composition in the brain changes with age. (**a** and **b**) CD45+ immune cell numbers of brain resident cells in male (**a**) and female (**b**) mice (n = 8/6 [4 weeks male/female]; n = 6/6 [adult male/female]; n = 4/3 [aged male/female]). (**c** and **d**) Quantification of each indicated immune cell in the brain from 4-week-old male (**c**) and female (**d**) mice (n = 8 in male mice and n = 6 in female mice). (**e** and **f**) Quantification of indicated immune cells in the brain from adult male (**e**) and female (**f**) mice (n = 6 in male mice and n = 6 in female mice). (**g** and **h**) Quantification of indicated immune cells in the brain of 24 months or older aged male (**g**) and female (**h**) mice (n = 4 in male mice and n = 3 in female mice). (**i** and **j**) Stacked percentages of each immune subsets on time points of their age 4 weeks, adult, aged were shown with connected lines. All n values refer to the number of mice used. *P<0.05, **P<0.01, and ***P < 0.001, calculated by One-way ANOVA (**i** and **j**) with Tukey’s post-hoc test (**a** and **b**). Data are shown as the mean +-SEM.

**Figure 3.**
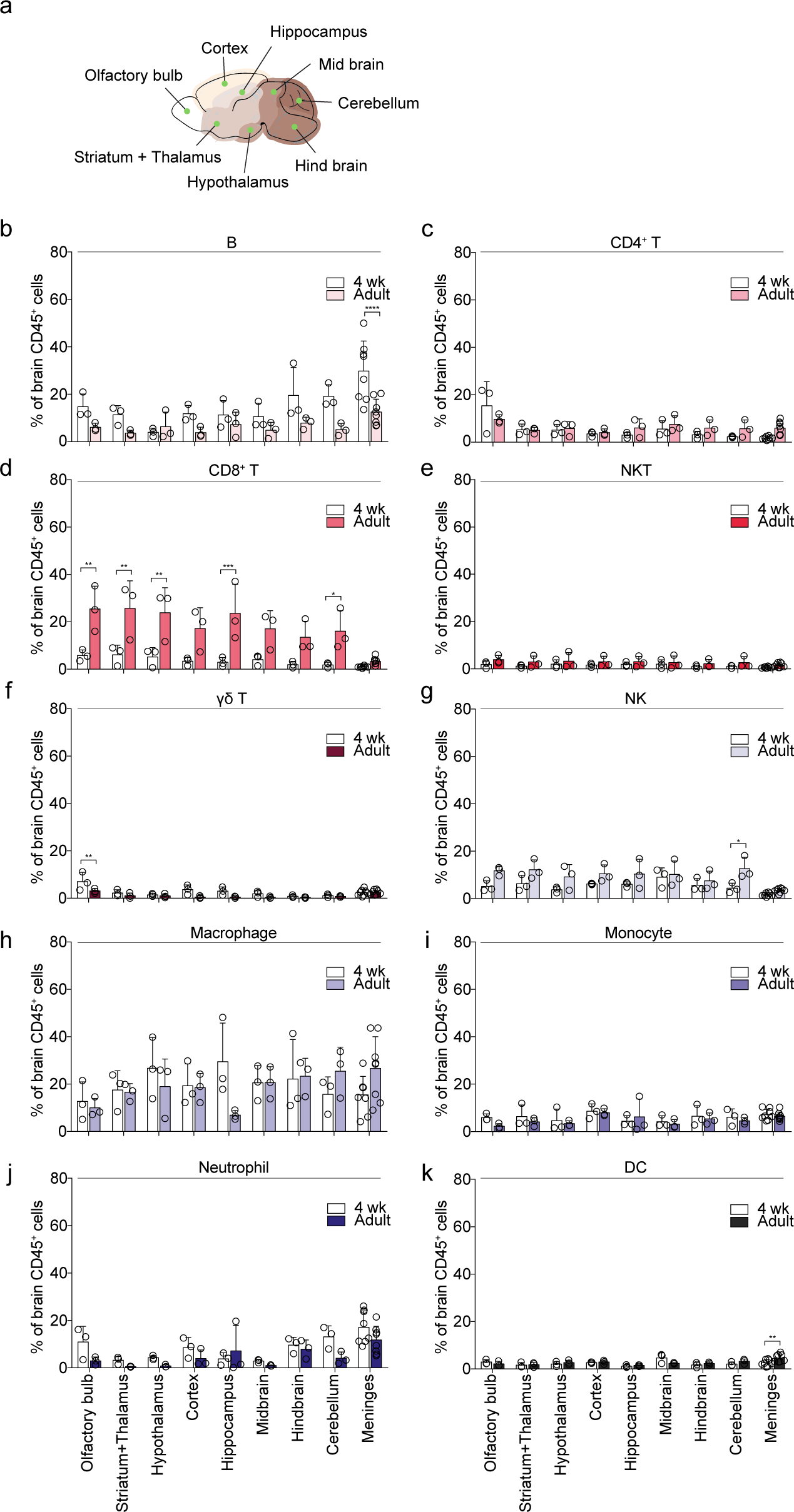
Immune cells exhibited no regional specificity. (**a**) Schematic representation in the sagittal section of the adult mouse brain with eight analyzed anatomical regions. (**b**-**k**) Quantification of (**b**) B cells, (**c**) CD4^+^ T cells, (**d**) CD8^+^ T cells, (**e**) NKT cells, (**f**) γδ T cells, (**g**) NK cells, (**h**) macrophages, (**i**) monocytes, (**j**) neutrophils, and (**k**) DC cells in each region of the brain from adult mice (n = 3 [olfactory bulb, striatum+thalamus, hypothalamus, hippocampus, cortex, midbrain, hindbrain and cerebellum] from three independent experiments. 4 or 5 mice brains were combined for each independent experiment; n = 8 [meninges]). All n values refer to the number of mice used. *P<0.05, **P<0.01, and ***P < 0.001, calculated by Two-way ANOVA with Sidak’s post-hoc test (**b**-**k**). Data are shown as the mean +-SEM.

### Brain-specific immune cells do not originate from maternal sources

Subsequently, we aimed to investigate the timing of immune cell colonization within the brain. Recent advances suggested that maternal microchimeric cells (MMc) convey health consequences for the offspring^9, 15–17^. The transfer of MMc from mother to fetus commences with maturing placentation, hence, with the onset of the second trimester in humans and around mid-gestation in mice^16^ and shapes neurodevelopment and behavior in mice^9^. However, whether these cells remain in the brain for the entire life has not been evaluated yet. To detect MMc in the brain, we crossed C57BL/6 female mice with BALB/c background CD45.1 congenic male mice to generate offspring with CD45.1/2^+^, H-2K^b/d^ and H-2D^b/d^ alleles (Fig 4a). Isolated immune cells were analyzed by FACS using the markers CD45.2 and CD45.1, after which CD45.2^hi^ cells were selected for further isolation. These cells were subsequently gated based on H-2K^b^/H-2D^b^ and H-2K^d^/H-2D^d^ markers, with only H-2K^b^/H-2D^bhi^ and CD45.2^hi^ cells being identified as MMc (Fig 4b). Intriguingly, we observed significantly fewer MMc in the brain compared to other tissues, which is a much smaller amount of MMc compared to those in the embryo brain (Fig 4c)^9^. Even though it is small, it contains immune cells including myeloid, B cells, and T cells (Fig 4d). This data suggests that the immune cells in brain parenchyma were born in the offspring and migrated into its brain, not from the mother, respectively.

**Figure 4.**
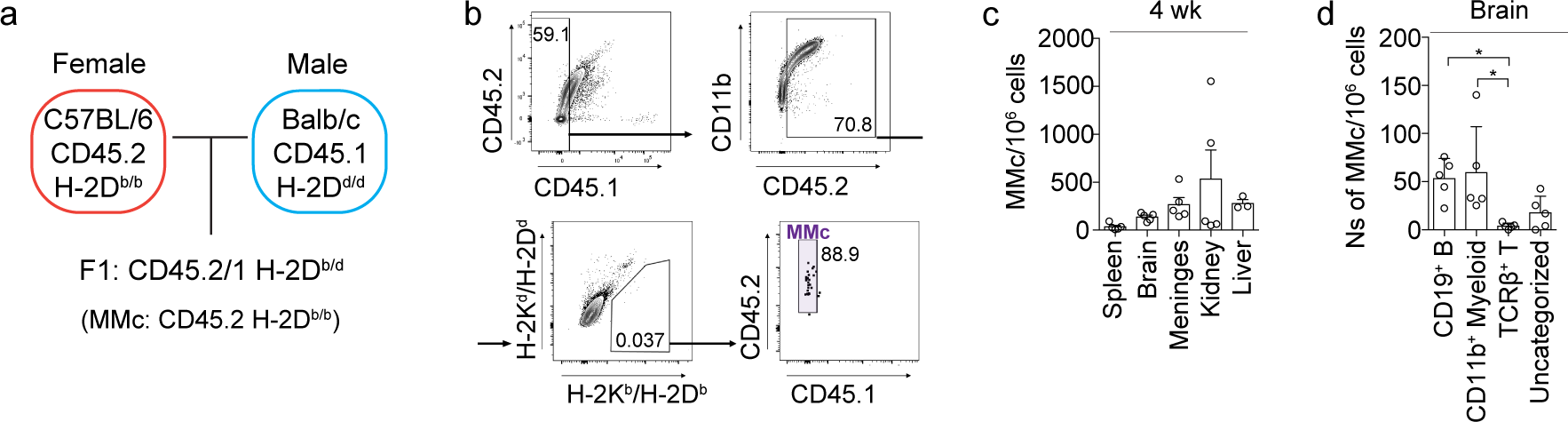
Brain-resident immune cells are not maternally derived. (**a**) Schematic diagram of the experimental method to detect MMc. Both H-2K and H-2D MHC I antigens, ptprc^a^ (CD45.1) were used for distinguishing MMC clearly. (**b**) Gating strategy for identification of maternal-derived immune cells. (**c**) Cell number of maternal-derived immune cells per 10^6^ cells in the spleen, brain, meninges, kidney, and liver (n = 5 [spleen, brain, meninges, and kidney], n = 3 [liver]). (**d**) Cell number of maternal-derived B, myeloid, and T cells per 10^6^ cells in the brain (n = 5). All n values refer to the number of mice used. *P<0.05, calculated by One-way ANOVA with Dunnett’s post-hoc test. Data are shown as the mean +-SEM.

### Brain-specific immune cells have different transcriptional profiling

If these cells originate within the offspring’s immune system, we inquired whether they exhibit unique signatures in comparison to immune cells in the blood. To gain deeper molecular insights into the immune cells residing within the brain, we isolated total immune cells and conducted sequencing analyses, alongside examining diluted microglia. And, the data was compared with data from blood immune cells. To minimize ex vivo activation and transcriptional activity during the isolation procedure, we generated single-cell suspensions under consistently cold conditions. We utilized a droplet-based RNA-seq approach and we pooled cells from five biological replicates. To identify transcriptionally distinct immune cells between blood and brain, we conducted dimensionality reduction and clustering through principal component analysis. This analysis revealed the overarching transcriptional differences between immune cells in the blood and those in the brain (Fig 5a). For a more detailed categorization, we further classified immune cells based on their marker genes into ten distinct subgroups, akin to the categories used in our FACS analysis (Fig 5b and c). Notably, immune cells within the brain parenchyma displayed heightened cytokine signaling, indicative of an activated immune response (Fig 5d and e). Particularly, in the ‘Cytokine Signaling in Immune System’ pathway, we observed the upregulation of 185 genes and downregulation of 40 genes, culminating in an overall augmentation of cytokines signaling within brain immune cells (Fig 5e). This consistent trend in cytokine signaling was observed across various subtypes of immune cells, with CD4+ and CD8+ T cells being particularly noteworthy (Extended Data Fig 3). Furthermore, the expression of key cytokines known to play pivotal roles in the brain, such as interferon (*Ifn*), tumor necrosis factor (*Tnf*), and chemokine ligands (*Ccls*), exhibited a substantial increase across all immune cell subsets when compared to blood immune cells (Fig 5f).

**Figure 5.**
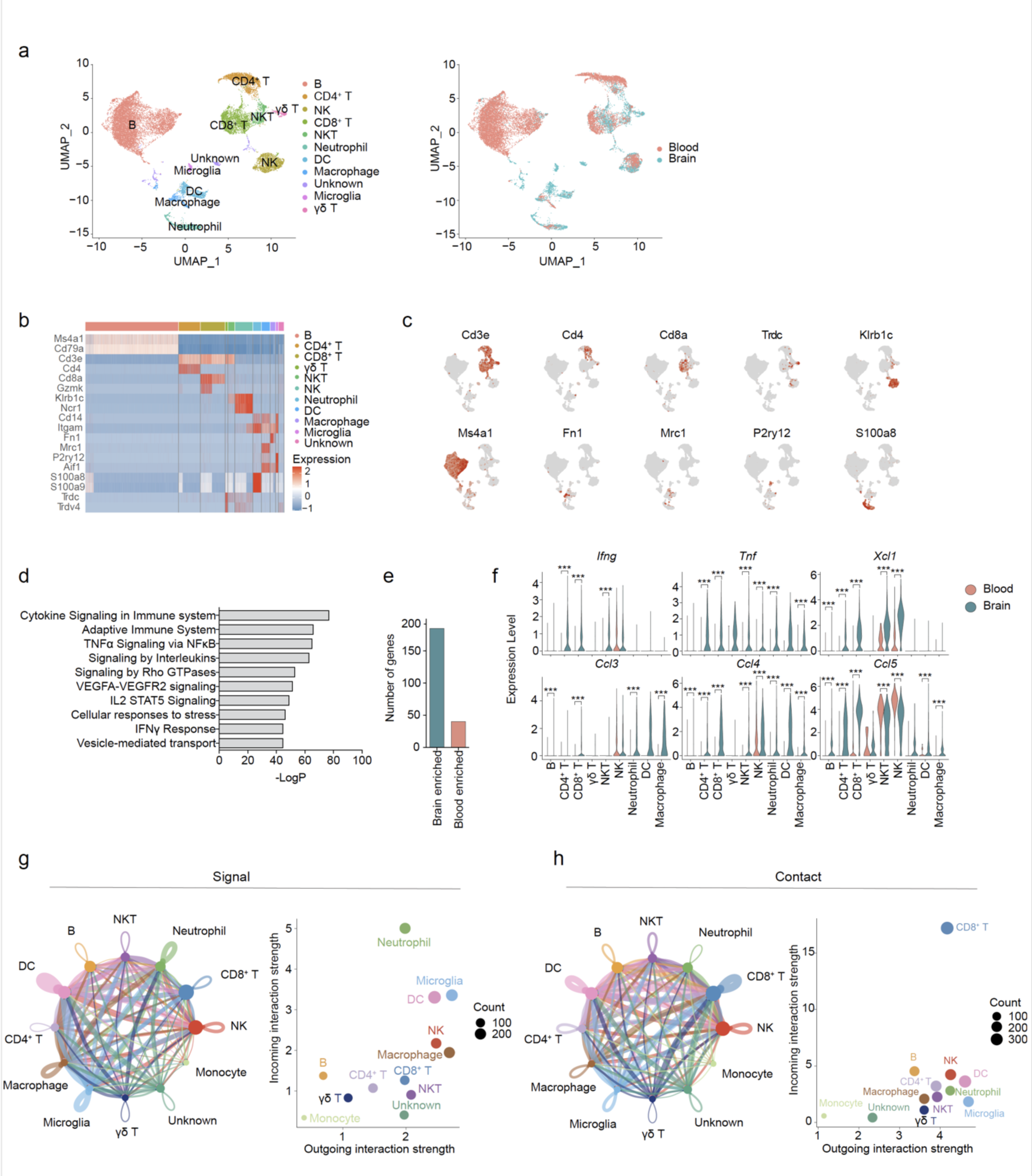
Brain-resident immune cells exhibit elevated cytokine expression. (**a**) UMAP embeddings in our data set. Colors represent cell type annotations (left), and origin tissues (right). (**b**) Heatmap demonstrating marker gene expressions for each annotated cell cluster. (**c**) UMAP embeddings colored by marker gene expression. Feature plots showing marker gene expression for each cluster in the UMAP. (**d**) Selected ten enriched biological pathway terms for genes that are up-regulated in the brain resident CD45^+^ immune cells than blood CD45^+^ immune cells. (**e**) Number of genes for Cytokine Signaling in the Immune system between total CD45^+^ brain resident immune cells and total CD45^+^ blood immune cells. (**f**) Violin plots showing indicated cytokine expression from each immune cell subsets in the brain and blood. (**g**) Circle plot of signaling networks of immune cells including microglia in the brain (left). Cell-cell interaction strengths plot indicating all cell types showing incoming and outgoing interactions (right). (**h**) Circle plot of direct contact networks of immune cells including microglia in the brain (left). Cell-cell interaction strengths plot indicating all cell types showing incoming and outgoing interactions (right). Statistical significance results for differentially expressed genes between cell types (**f**) were calculated by Wilcoxon rank-sum test with Bonferroni correction. For B cells: *Ccl4,* p=5.65e-205; *Ccl3,* p=9.30e-151; *Xcl1,* p=5.62e-127; *Ccl5,* p=6.43e-73. For CD4+ T cells: *Ifng,* p=1.07e-130; *Ccl5,* p=2.42e-122; *Xcl1,* p=3.41e-116; *Tnf,* p=2.69e-111; *Ccl4,* p=2.67e-104. For CD8+ T cells: *Ccl5,* p=1.86e-231; *Xcl1,* p=3.99e-228; *Ccl4,* p=2.31e-216; *Tnf,* p=5.10e-131; *Ifng,* p=4.67e-129; *Ccl3,* p=1.43e-70. For NK cells: *Xcl1,* p=1.45e-136; *Ccl5,* p=8.73e-122; *Tnf,* p=1.51e-37; *Ccl4,* p=1.90e-12. For NKT cells: *Xcl1,* p=9.85e-21; *Ccl5,* p=2.02e-12; *Ccl4,* p=2.41e-08; *Ifng,* p=1.21e-04; *Tnf,* p=2.81e-04. For DC: *Ccl4,* p=6.23e-03. For Macrophages: *Ccl3,* p=1.82e-33; *Ccl4,* p=2.46e-15; *Tnf,* p=1.10e-08; *Ccl5,* p=2.21e-04. For Neutrophils: *Tnf,* p=3.16e-10; *Ccl3,* p=1.28e-06; *Ccl4,* p=1.85e-04.

Given the conspicuous upregulation of cytokine expression, we hypothesized that these cytokines might have the potential to influence cells within the brain. It is now well-established that certain levels of cytokine are necessary to maintain neuronal homeostasis. To delve deeper into this hypothesis, we examined interactions involving cytokines, signaling, and cell-to-cell contact across various immune cells, including microglia, employing the CellChat tool^18^. CellChat assesses the strength of cell-to-cell interactions in scRNA-seq data using a ligand-receptor interactions database. The results revealed that microglia, a pivotal component in maintaining brain homeostasis, engaged in multifaceted interactions with various immune cells, both in terms of ‘signaling’ and ‘direct contact’ (Figure 5g and h). Particularly noteworthy was the robust interaction observed between microglia and CD8^+^ T cells, especially in the context of ‘direct contact’ (Fig 5h and Extended Data Fig 4). Overall, our findings suggest that immune cells residing within brain parenchyma exhibit a heightened state of activation compared to blood immune cells. Furthermore, these cells engage in intricate interactions, particularly with microglia, and may play a significant role in influencing brain function through interaction with microglia.

### AD mice exhibited new signaling interactions between microglia and immune cells

Our findings demonstrated that immune cells reside in the healthy brain, and their composition and characteristics differ from immune cells residing in other organs. Additionally, through analysis using CellChat, we observed robust interactions among immune cells and strong interactions with microglia. Notably, a prominent feature of brain immune cells is the strong expression of several cytokines. However, the expression of cytokines is often linked to their detrimental effects on brain-resident cells, associated with neuronal pathophysiology, and implicated in the progression of the disease^6, 19, 20^. To explore the distinctions in induced cytokine expression within the healthy brain under pathological conditions, we investigated brain immune cells in Alzheimer’s disease (AD), which is a neurodegenerative brain disorder. We conducted a comparative analysis of two distinct AD mouse models, derived from human APOE4 expressing mouse (E4), namely the amyloid-β deposition mouse model, APP/PS1-21(A/PE4), and the tauopathy model (TE4) (Fig 6a)^21^. However, unexpectedly, the expression levels of *Ifn* and *Tnf*, which are commonly suggested as key molecules inducing AD pathology, did not show significant differences between the AD mouse model and the control E4 mice (Extended Data Fig 5a). This observation suggests that the cytokines themselves may not be the primary contributor to the induction of neuropathological changes in the brain. In line with prior research, a quantitative assessment of the proportion of CD45 high brain immune cells in these mouse models revealed deviations from the observed immune cell proportions in the control E4 mice. Particularly, the tauopathy model exhibited a significant increase in the CD8^+^ T cell population (Fig 6b). Subsequently, we delved into the possibility of novel underlying mechanisms in cell-to-cell communication. In CellChat analysis, we found that contact signals were relatively stable in both the AD mouse and control E4 mouse (Extended Data Fig 5a). Conversely, a paradigm shift in the signaling environment of microglia was evident in the AD model. Signaling pathways that occur under normal conditions were dampened in the AD condition, and new ligand-receptor interactions that were absent under normal conditions emerged, which include *Igf1-Igf1r, Csf1-Csf1r, Ccl9-Ccr1,* and *Ccl2-Ccr2* (Fig 6c and d). Additionally, we observed an increase in *Vegfb* and *Spp1* signaling in the TE4 mouse model. Taken together, these findings offer us two significant insights: 1. The cytokine expression of immune cells in the healthy brain and the AD-affected brain shows no significant differences. 2. Nevertheless, new ligand-receptor interactions or an increased number of cytotoxic immune cells might be emerging mechanisms contributing to AD conditions. This implies that the emergence of new interactions between microglia and brain immune cells in the AD mouse model underscores a comprehensive interplay that collectively contributes to the differences in both qualitative and quantitative characteristics of brain immune cells.

**Figure 6.**
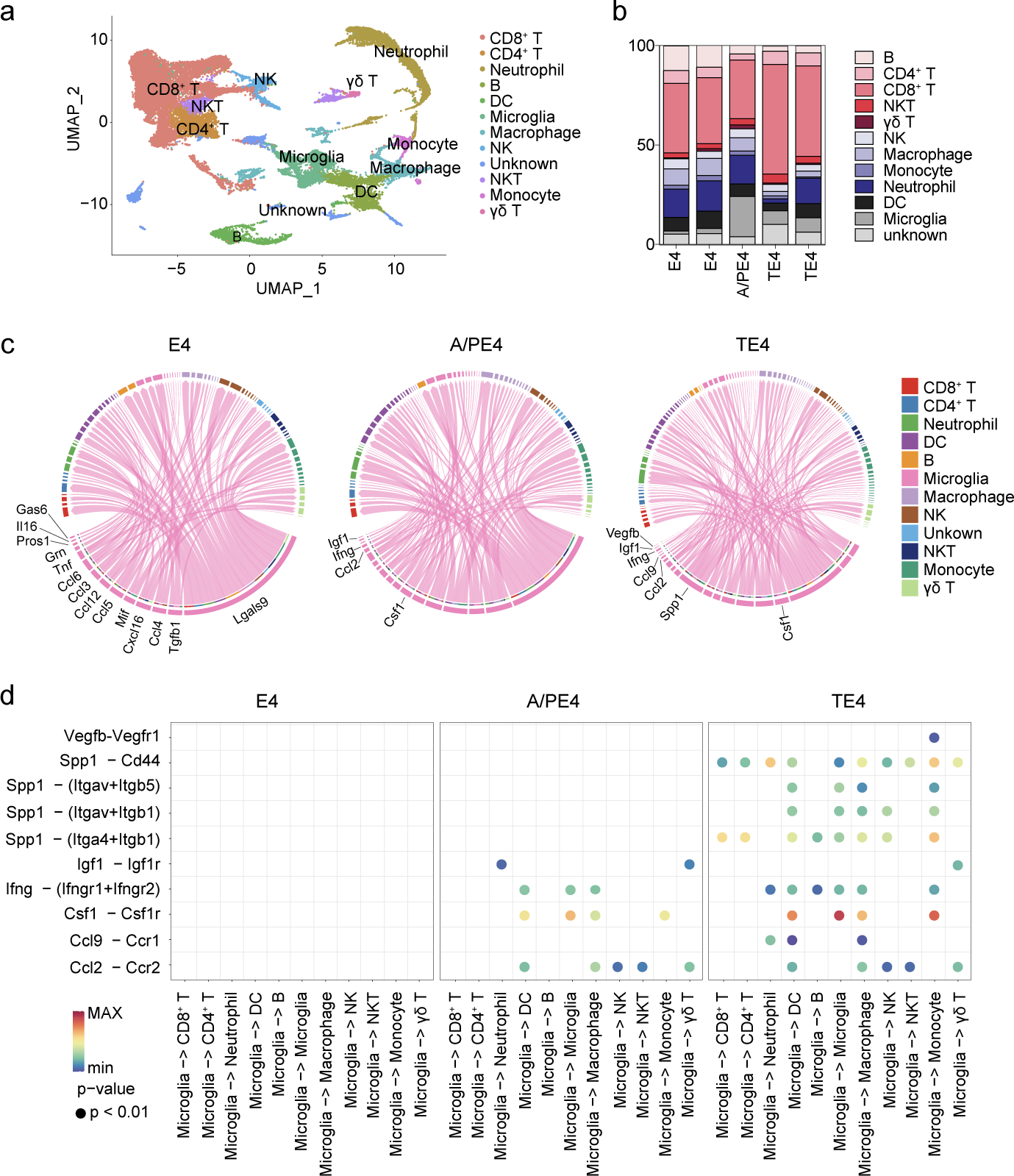
Signal interactions between brain-resident immune cells and microglia are perturbed in an Alzheimer’s disease model. (**a** and **b**) UMAP plot (**a**) and bar graphs (**b**) representing immune cell clusters of the integrated dataset in all brain CD45 high samples from E4, A/PE4, and TE4 mice. (**c**) Chord diagram showing each ligand-receptor pattern and their weights from microglia cells to all indicated cells. (**d**) Dot plots indicating signaling molecules between microglia and each immune cell in E4, A/PE3, and TE4 mice.

## Discussion

Our study has significantly transformed our understanding of immune cell presence within the brain. Contrary to the traditional concept of immune privilege, we have provided compelling evidence supporting the existence of immune cells in the brain parenchyma, akin to their presence in another non-lymphoid organs^3–5^. This finding encompasses ten major immune cell types, including macrophage, monocytes, dendritic cells, neutrophils, NK cells, B cells, CD4^+^ T cells, CD8^+^ T cells, γδ T cells, and NKT cells, which exhibit varying degrees of abundance throughout an individual’s lifespan. These observations yield three significant insights. 1. The Brain shares commonalities with peripheral non-lymphoid organs by harboring resident immune cells. 2. The presence of immune cells in the brain does not necessarily signify a previous infection. 3. Brain-resident immune cells do not originate from maternal sources during embryogenesis. This paradigm shift suggests that these brain-specific immune cells may play specialized roles in maintaining brain homeostasis and responding to unique brain-related challenges. The presence of immune cells in the brain parenchyma raises intriguing questions about their roles in maintaining brain homeostasis and function. While their numbers in the brain parenchyma may be relatively sparse compared to other tissues, their localization within the brain suggests potential specialized functions. Further investigations should focus on uncovering their specific functions and contributions to brain health, as well as the mechanisms behind their recruitment and maintenance.

Additionally, our study has uncovered a dynamic landscape of immune cell interactions and cytokine signaling within the brain. For an extended period, research in the field predominantly focused on the detrimental effects of cytokines and the presence of immune cells in the brain on brain-resident cells, such as neurons, microglia, and astrocytes. For example, infiltration of CD8+ T cells is significantly enhanced in disease-associated brain areas in neurodegenerative diseases such as multiple sclerosis (MS), AD, and Parkinson’s disease (PD). CD8^+^ T cells produce inflammatory cytokines IFN-γ and TNF, which activate other immune cells, including microglia, thus enhancing the inflammatory microenvironment^19^. Additionally, these induced cytokines have also been implicated in neuronal death and impaired motor function in murine models of PD and AD, and are elevated in the blood of patients^20,21^. However, our findings align with recent advancements that have begun to shed light on the essential roles of cytokines in maintaining brain homeostasis and ensuring proper development, extending beyond their effects on pathological conditions. Over the last two decades, an increasing number of studies have investigated the effects of cytokines on brain development, social behavior, learning and memory, and emotion under physiological conditions^22,23^. A major limitation of previous research is the lack of specificity regarding the source of cytokines. Our new findings might prompt others to consider the brain-resident immune cells as a primary source of cytokines in physiological conditions. Surprisingly, when we examined cytokine expression levels in each immune cell from the AD model, we observed no significant changes compared to the WT mice, especially for IFN-γ and TNF. Intriguingly, this model exhibited the emergence of new signaling pathways related to Vegfb, Spp1, Igf1, Csf1, Ccl2, and Ccl9. These findings suggest that while the healthy brain employs cytokines as essential mediators of normal function, the AD-affected brain may adapt by invoking alternative signaling cascades or recruiting more cytotoxic immune cells. Such adaptations underscore the remarkable plasticity of the brain’s immune response mechanisms and warrant further investigation into the complex interplay between cytokines and brain health.

In summary, our study underscores the emerging significance of neuroimmunology in shaping our understanding of brain function and offers promising prospects for insights into normal brain development and AD pathogenesis.

## Materials and Methods

### Mice

Wild-type C57BL/6J mice were purchased from Jackson Laboratory and maintained under specific pathogen-free conditions at the University of Pennsylvania animal facility. Three- to ten-months-old mice were used as the “Adult” group. Mice more than twenty-four months old were used as the “Aged” group. The BALB/C background CD45.1 male mice were purchased from the Jackson Laboratory (JAX #006584) and bred with a C57BL/6J female for detecting MMc. All experiments and breeding were done according to the guidelines of the Institutional Animal Care and Use Committee (IACUC) of the University of Pennsylvania.

### Immune cell preparation

To stain the vascular circulating immune cells, we injected 3 μg of anti-mouse CD45 (Clone:30-F11) in 300 μl of PBS via the tail vein (i.v.). Three minuts Later, the mice were perfused with 25 ml of cold PBS under isoflurane anesthesia. We collected a whole mount of dural meninges from the mouse skullcap, which was detached from the skull, using sharp forcepts^24^. The whole brain was then removed from the skull. For the dissection of the mouse brain, we anatomically dissected the following regions: 1) olfactory bulb, 2) striatum and thalamus, 3) hypothalamus, 4) hippocampus, 5) cortex, 6) midbrain, 7) hindbrain (Pons and Medulla oblongata), and 8) cerebellum. We collected at least four mouse brains for the brain dissection experiment. Lateral, 3rd, and 4th choroid plexus were collected separately during the anatomical dissection^25^. The whole brain or dissected brain were chopped in 1.5 ml tube with 400 μl of 10% FBS contained RPMI1640 media and transferred into glass vial tubes containing digestion solution consisting of 10% FBS, 2 μg/ml Collagenase IV (Sigma), and 50 μg/ml DNase I in RPMI1640 media. The samples were stirred for 25 minuts at 37°C with a cross-shaped magnetic bar, washed twice, and then centrifuged through a 40% percoll layer on top of a 75% percoll solution for 20 minutes at room temperature, without interruption. After removing the upper myelin layer, mononuclear cells were collected and washed twice for further experiments.

### Flow cytometry

Single suspension cells were stained using LIVE/DEAD™ Fixable Near-IR Dead Cell Stain Kit (Thermo Fisher Scientific) to exclude dead cells. Surface antigens were stained for 30 min at 4°C with indicated antibodies (Table.1) after Fc blocking for 4°C 30 min with anti-mouse CD16/32 (BD) without wash. After washing with PBS, samples were analyzed on BD FACSymphony™ A3. Data was collected through FACSDiva (BD Pharmingen). Analysis was done with FlowJo software (BD Biosciences).

### Cell sorting and scRNA-Seq library construction

Three-month-old male C57BL/6 mice (n = 5) were transcranial perfused with ice-cold PBS after staining the vascular circulating immune cells as described. Blood samples were collected by heart puncture before perfusion. Blood samples underwent red blood cell lysis for 4 min at room temperature. Brains were physically chopped before undergoing digestion with Collagenase VI and DNase I and underwent percoll gradient centrifugation as described. Obtained single-cell suspension from the brain and blood was stained as described. Stained cells were washed and FACS sorted by BD FACS Aria II for isolating live blood CD45^+^ and brain resident CD45^+^ cells (Purity 98%<). After washing once, FACS-sorted cells were encapsulated in a 10× Chromium instrument, and libraries were constructed using a Single Cell 3′ Reagent Kit (10X Genomics V3 Chemistry) following the manufacturer’s instructions. Subsequently, the libraries were sequenced using the NovaSeq 6000 platform.

### scRNA-seq computational analysis

The initial processing of scRNA-seq data was performed using the Cell Ranger pipeline (v.7.0.0), which was downloaded from 10X Genomics (https://support.10xgenomics.com/single-cell-gene-expression/software/downloads/7.0). Base call files were demultiplexed into FASTQ files using the cellranger mkfastq with Illumina bcl2fastq2 software (2.20.0.422). The resulting reads were aligned to the mm10 transcriptome (refdata-gex-mm10-2020-A) using cellranger count^26^. For subsequent analysis, we employed R (4.2). The Cell Ranger output is organized using Seurat (v.4.0.6)^27^. Different cut-offs were set for two distinct datasets. In the brain dataset, cells with fewer than 300 unique genes or more than 7,300 genes per cell were filtered out. Additionally, cells with a mitochondrial gene percentage greater than 5% or a percentage of the largest gene greater than 15% were excluded. In the blood dataset, cells with fewer than 350 unique genes, more than 6,000 genes per cell, a mitochondrial gene percentage greater than 7%, or a percentage of the largest gene greater than 10% were removed. DoubletFinder (2.0.3) was employed to eliminate doublets^28^. For the brain dataset, the parameters used were pN= 0.2, pK=0.11, and dp=0.055, while for the blood dataset, pN=0.15, pK=0.08, and dp=0.075 were applied.

After the quality control process, the blood and brain datasets were merged using the IntegrateData function. The SelectIntegrationFeatures, PrepSCTIntegration, and FindIntegrationAnchors functions were utilized to identify the common anchor sets used in the IntegrateData function. Following data merging, the RunPCA function was applied to compute principal components for all genes. The first 30 components were selected for UMAP and Shared Nearest Neighbor clustering. The Louvain algorithm was optimized with a resolution of 1.0 and applied to the shared nearest neighbor graph using the FindClusters function. The annotation of clusters was conducted based on the presence of common cell type markers, resulting in the identification of 10 distinct cell types: B cells (Cd79a, Ms4a1), gamma delta T cells (Cd3e, Trdc), CD4^+^ T cells (Cd3e, Cd4), CD8^+^ T cells (Cd3e, Cd8a), NKT cells (Cd3e, Klrb1c), NK cells (Klrb1c, Ncr1), Macrophages (Fn1), Dendritic Cells (Cd14, Mrc1), Neutrophils (S100a8, S100a9), and Microglia (P2ry12, Aif1).

### Differential expression analysis

For all differential expression testing, we employed the Wilcoxon rank-sum test with Bonferroni correction using the FindMarkers function. Genes were considered significant if they exhibited a minimum log2 fold change greater than 0.1 and were expressed in a minimum of 10% of cells within the cluster of interest.

Metascape (www.metascape.org) was used for biological pathway analysis (29). Specifically, we utilized BioCarta Gene Sets, Canonical Pathways, Hallmark Gene Sets, Reactome Gene Sets, KEGG Pathway, and WikiPathways.

### Cell-cell communication analysis

We reconstructed cell–cell interactions based on the expression of known ligand–receptor pairs in different cell types using the CellChat (1.6.1)^29^. We selected the “Secreted Signaling” and “Cell-Cell Contact” databases curated by CellChat. Preprocessing functions, namely identifyOverExpressedGenes and identifyOverExpressedInteractions with default parameters, were applied to datasets. Additionally, we utilized the functions computeCommunProb, computeCommunProbPathway, and aggregateNet with default parameters for the analyses.

### Processing and analysis of public scRNA-seq datasets

GSE221856 scRNA-seq dataset that contains Brain CD45 high cells (GSM6900873, GSM6900875, GSM6900876, GSM6900878, GSM6900880) are used for analysis^21^.

## Data availability

All sequencing data will be made publicly available upon publication.

## Acknowledgements

We thank all members of the Yim laboratory for helpful discussions and critical analysis of the manuscript.

## Funding

This publication was made possible by P30 ES013508 from the National Institute of Environmental Health Sciences, NIH. Its contents are solely the responsibility of the authors and do not necessarily represent the official views of NIEHS, NIH.

## Author contributions

S.H., J.J., K.P., and Y.S.Y. designed the experiments and/or provided advice and technical expertise. S.H. and K.P. performed the experiments. S.H., J.J., K.P., and Y.S.Y. wrote the manuscript.

## Competing interests

The authors declare no competing interests.

---

**Extended Data Figure 1.**
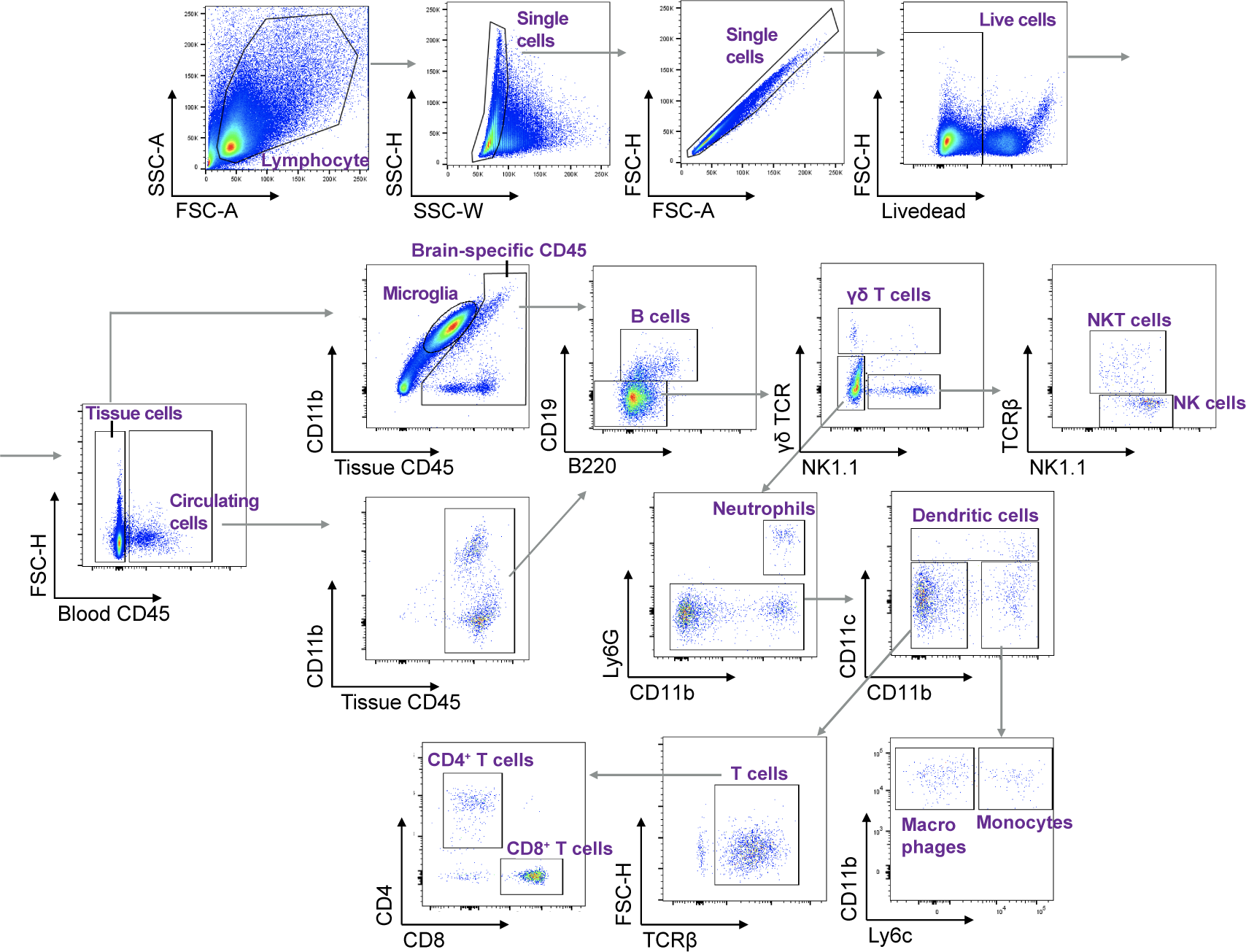
Gating strategy for multi-color flow cytometry analysis of each immune cell type in the brain.

**Extended Data Figure 2.**
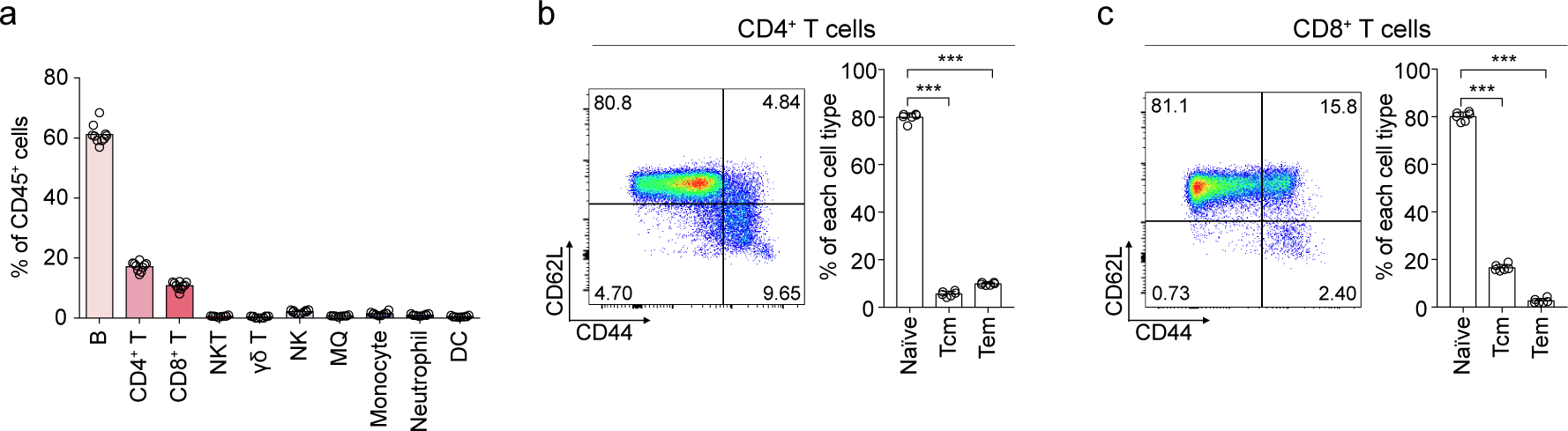
Spleen immune cell profile (**a**) and CD62L/CD44 profile of splenic CD4+ and CD8+ T cells (**b** and **c**) for showing the non-infected condition of mice used in this paper. (**a**) Quantification of indicated immune cells in spleen from adult mice. (b-c) Representative flow cytometric plot of naive (CD62L^+^CD44^−^), central memory (Tcm: CD62L^+^CD44^+^), and effector memory (Tem: CD62L^−^CD44^+^) in splenic CD4^+^ T cells (**b**) or CD8^+^ T cells (**c**) (left). Quantification of naïve, Tcm, and Tem in splenic CD4^+^ T cells (**b**) or CD8^+^ T cells (**c**) as indicated (right). All n values refer to the number of mice used. ***P < 0.001, calculated by One-way ANOVA with Dunnett’s post-hoc test (**b** and **c**). Data are shown as the mean +-SEM.

**Extended Data Figure 3.**
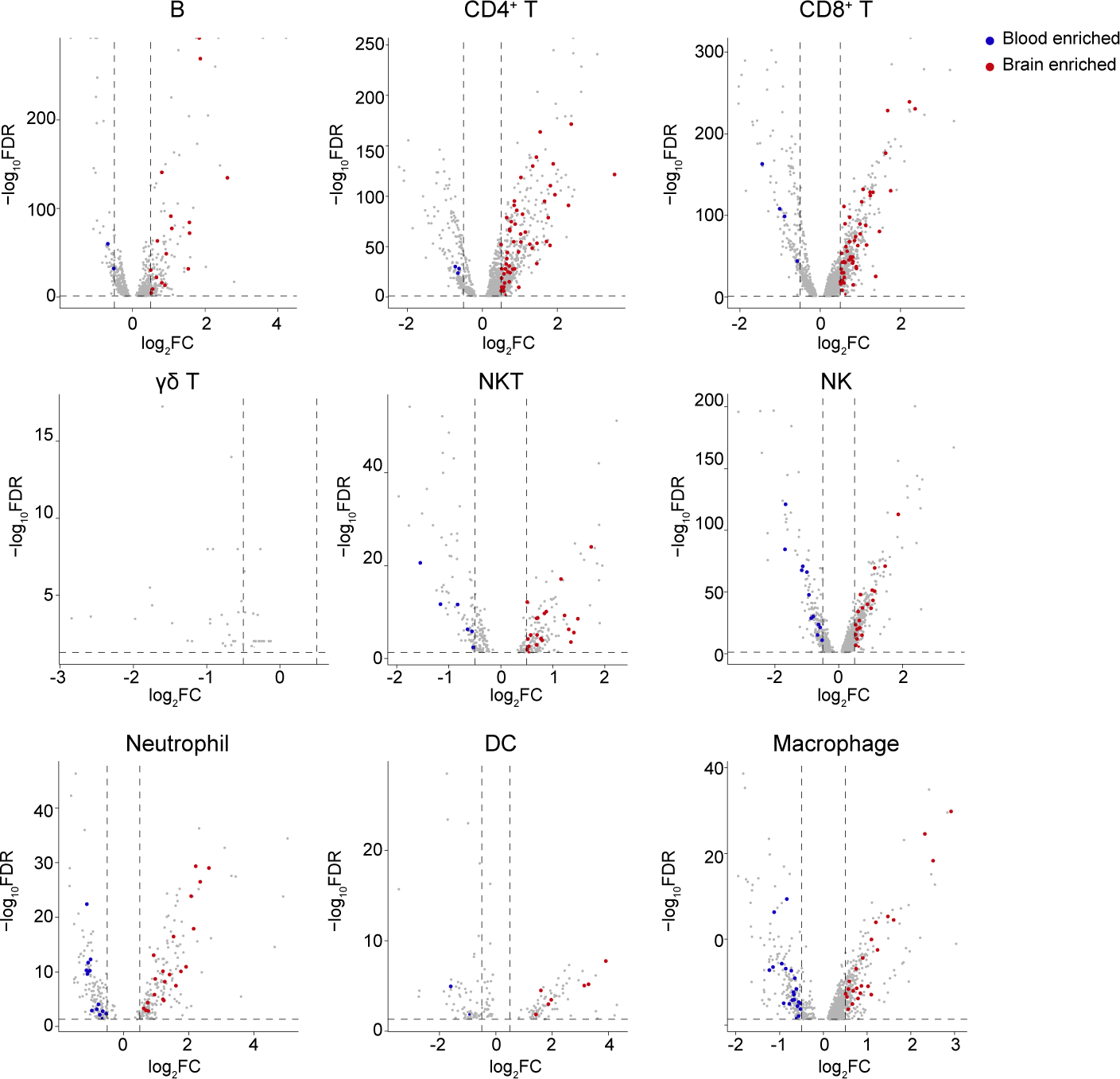
Brain immune cell subtypes are highly enriched in genes related to cytokine signaling compared to blood immune cell subtypes. Volcano plots highlighting genes that are differentially expressed (false discovery rate (FDR) < 0.05 and log2 fold change > 0.5) between brain immune cell subtypes and control blood immune cell subtypes (gray). For genes in the “Reactome cytokine signaling of the immune system” pathway, those enriched in blood immune cell subtypes are highlighted in blue, and those enriched in brain immune cell subtypes are highlighted in red. Cell subtypes include B cells, CD4^+^ T cells, CD8^+^ T cells, γδ T cells, NKT cells, NK cells, Neutrophils, DCs, and Macrophages (Upper left to lower right).

**Extended Data Figure 4.**
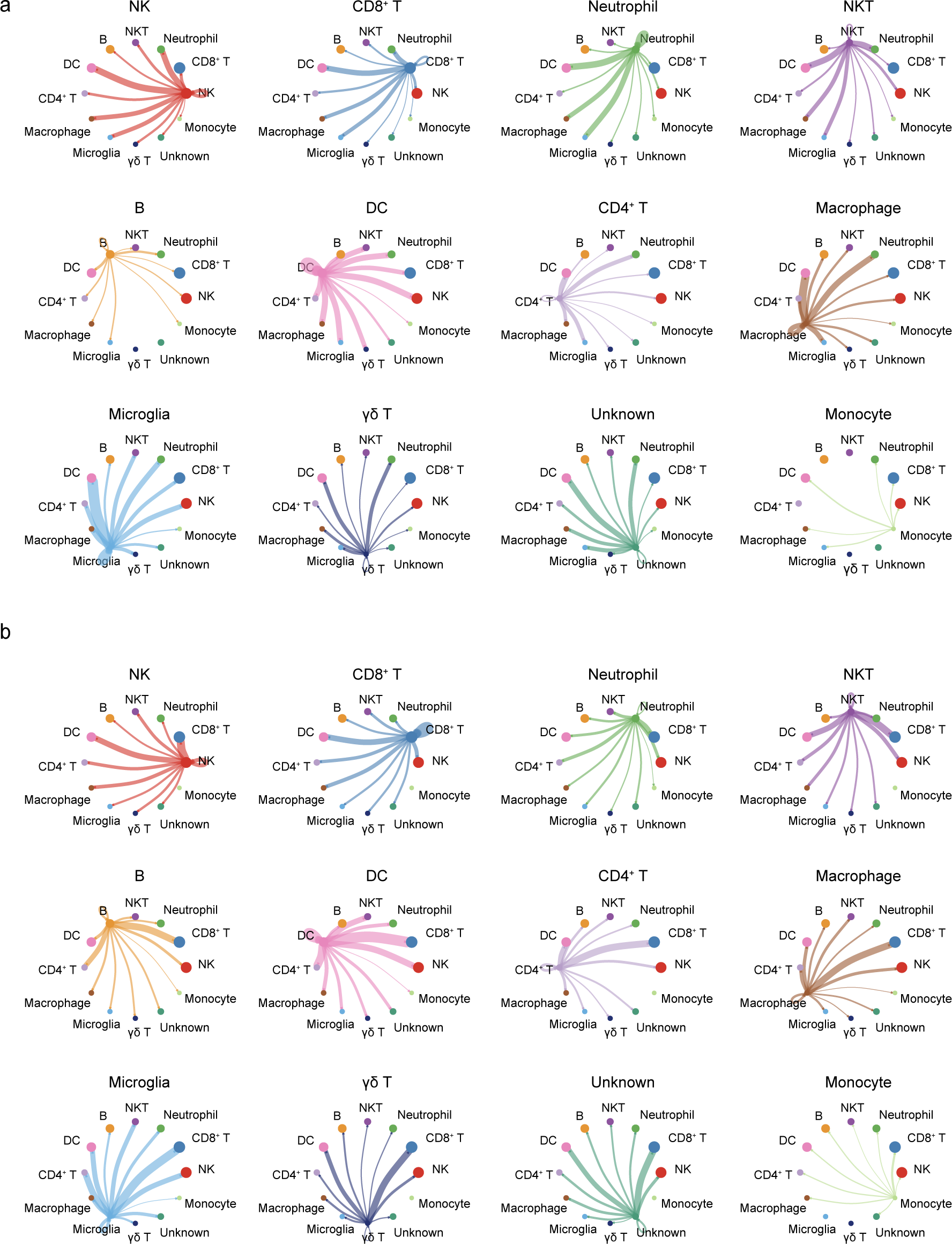
Cell-to-cell communications across the brain immune cell subtypes. Circle plots of interaction network of (**a**) “Secreted Signaling” and (**b**) “Cell-Cell Contact”. Cell subtypes include NK cells, CD8^+^ T cells, Neutrophils, NKT cells, B cells, DCs, CD4^+^ T cells, Macrophages, Microglia, γδ T cells, Unknowns, Monocytes (Upper left to lower right).

**Extended Data Figure 5.**
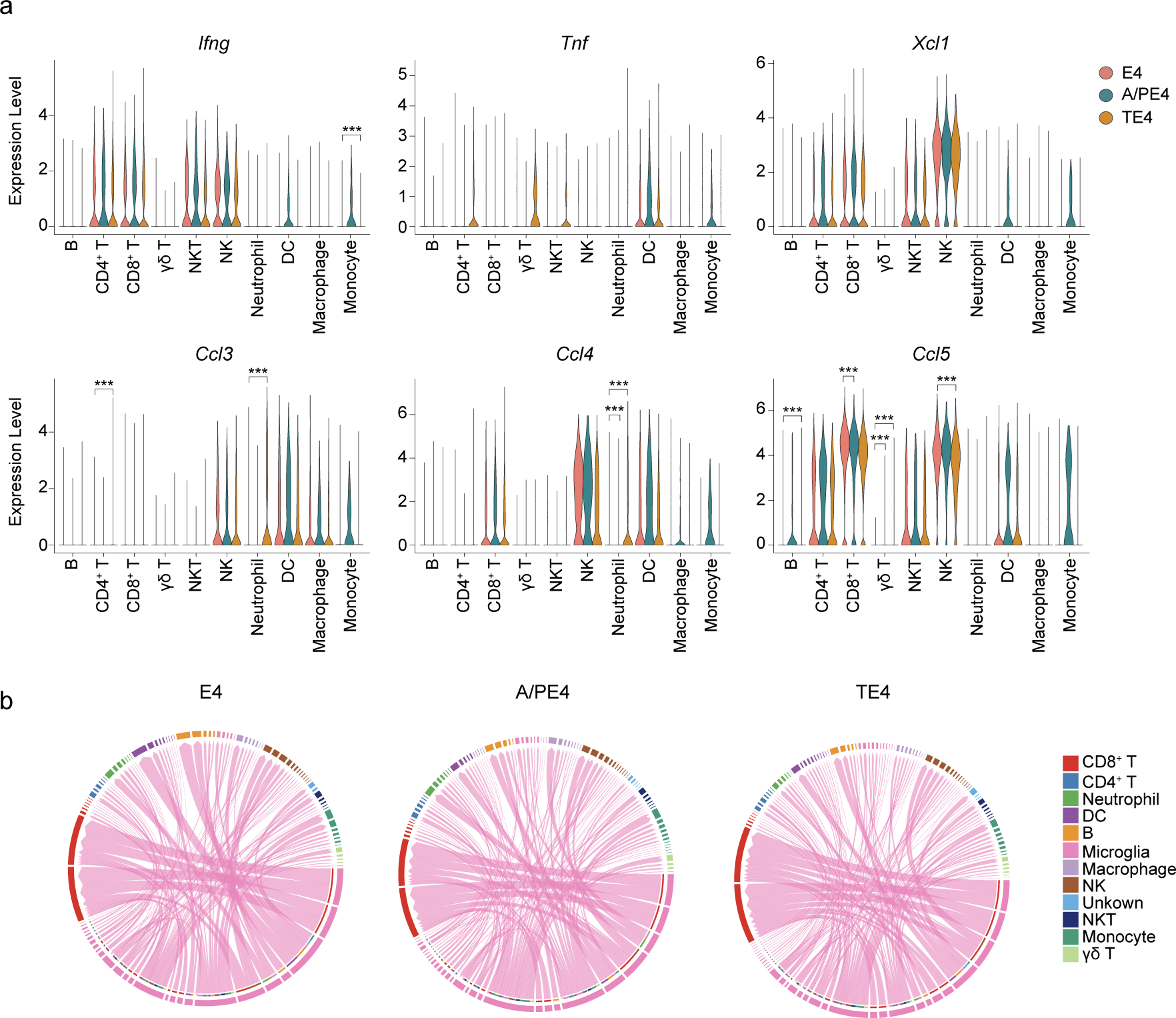
(**a**) Violin plots showing the expression *Ifng*, *Tnf*, *Xcl1*, *Ccl3*, *Ccl4*, and *Ccl5* in the immune cells from the control E4 (red), A/PE4 (blue), and TE4 mice (yellow). (**b**) Chord diagrams of control E4 (left), A/PE4 (middle), and TE4 mice (right) showing each ligand (microglia)-receptor (other peripheral brain immune cells) patterns and their weights. Statistical significance results for differentially expressed genes between cell types (**a**) were calculated by Wilcoxon rank-sum test with Bonferroni correction. 1) A/PE4 vs E4: CD8+ T cells; Ccl5, p=8.20e-24. γδ T cells; Ccl5, p=5.11e-07. Neutrophils; Ccl4, p=6.44e-70. 2) TE4 vs E4: B cells; *Ccl5,* p=1.73e-10. CD4+ T cells; *Ccl3,* p=9.80e-32. γδ T cells; *Ccl5,* p=1.51e-04. NK cells; *Ccl5,* p=5.77e-04. Monocytes; *Ifng,* p=4.41e-07. Neutrophils; *Ccl3,* p=3.41e-05, *Ccl4,* p=2.58e-24.

## Notes

### Competing Interest Statement

The authors have declared no competing interest.

